# Crystal Structure of the putative tail fiber protein gp53 from the *Acinetobacter baumannii* bacteriophage AP22

**DOI:** 10.1101/518761

**Authors:** Lada V. Sycheva, Mikhail M. Shneider, Anastasia V. Popova, Rustam H. Ziganshin, Nikolay V. Volozhantsev, Konstantin A. Miroshnikov, Petr G. Leiman

## Abstract

This report describes the structure of a putative tail fiber protein of the *Acinetobacter baumannii* bacteriophage AP22. The target host range of strictly lytic bacteriophage AP22 includes many clinical isolates of *A. baumannii* from hospitals in Chelyabinsk, Nizhny Novgorod, Moscow and St. Petersburg (Russia), but its host cell binding apparatus remains uncharacterized. Here, we report the crystal structure of the C-terminal fragment of AP22 gene product 53 (gp53) one of its two putative host cell-binding proteins. We show that gp53 forms a trimeric fiber and binds ethylene glycol and glycerol molecules that represent known surrogates of the oligosaccharide backbone. However, despite its structural similarities to other phage/virus host cell-binding fibers and its binding to small sugar-like molecules, gp53 did not inhibit AP22 infection and its role in the infection process remains unclear.

## Introduction

*A. baumannii* is a Gram-negative nonfermenting aerobic bacterium that frequently causes hospital-acquired infections world-wide because of its resistance to multiple drug classes [1,2]. The major types of infections caused by *A. baumannii* include pneumonia, bacteremia, endocarditis, skin and soft tissue infections, and meningitis [3]. *A. baumannii* displays a high capacity to acquiring new genetic elements that determine its antibiotic resistance leading to the persistence of this microorganism in the hospital environment and emergence of pandrug-resistant strains [4,5]. This property makes *A. baumannii* infections difficult to treat with the available antibiotics and alternative treatment strategies are being explored. One of them is the use of bacteriophages or their component proteins [6].

Strictly lytic *A. baumannii* phage AP22 was isolated from clinical materials and classified as a member of *Myoviridae* family [7]. AP22 was found to lyse 68% of 130 tested clinical multidrug-resistant *A. baumannii* isolates from various Russian clinics. AP22 therefore represents a promising anti-*A. baumannii* tool [7]. Its biological properties and infection parameters were characterized [3] and its lytic cycle was visualized with the help of atomic force microscopy [8]. However, the interaction of AP22 with its bacterial host on the receptor level remains unclear.

We performed bioinformatic analysis of AP22 virion proteins and identified two putative receptor binding proteins (RBPs), encoded by genes 53 and 54 (gp53 and gp54). The presence of several RBPs on the phage particle is often associated with expanded host cell range [9]. On the other hand, many phages carry multiple RBPs but do not show particularly broad host range. Bacteriophage T4 with its long and short tail fibers that are responsible for the initial host recognition and irreversible cell surface binding, respectively, is perhaps the best example of such functional specialization [10].

In this work, we analyzed the functional role of two putative AP22 RBPs gp53 and gp54 in infection. We found that the presence of gp54 in the cell culture medium very strongly inhibits AP22 infection, but gp53 produces no detectable effect. We also report the crystal structure of the C-terminal fragment of gp53, gp53C. We found that gp53C consists of a globular lectin-like C-terminal head domain and a slender midsection, an organization that is typical to host-cell binding fibrous proteins found in other phages and viruses (e.g. phage T7 gp17 [11] and T4 gp12 [12]) Furthermore, we show that the C-terminal lectin-like domain of gp53 binds ethylene glycol and glycerol molecules. These findings contradict the functional role of gp53 as determined in the infection assay and suggest that gp53 might be important for host cell binding in some yet not identified conditions.

## Materials and methods

### Cloning, expression and purification

Gene 53 of bacteriophage AP22 was PCR-amplified and cloned into the pEE3 vector. pEE3 was derived from pET23a (Novagen) by insertion of the sequence encoding His(6)-tag with TEV protease digestion site 8 nucleotides downstream from ribosome-binding site replacing the T7 tag. Protein expression was performed in *E. coli* B834(DE3) cells grown in 1 L of the 2TY medium supplemented with 200 μg/ml ampicillin at 37°C. When the cell culture reached an optical density of 0.6 (measured at the wavelength of 600 nm), it was cooled down to 18°C and gp53 expression was induced by addition of isopropyl-β-D-thiogalactoside to a final concentration 1 mM. The expression continued overnight. Cells were harvested on the following day by centrifugation at 8000g, 4°C for 10 min. The cell pellet was resuspended in 50 mM Tris pH 8.0, 300 mM NaCl, 5 mM imidazole. Cells were lysed by sonication. Lysate was centrifuged at 25000g, 4°C for 10 min. The supernatant was applied to a Ni^2+^ column (5 ml GE HisTrap FF Crude). The non-specifically bound material was washed away with 10 column volumes of 50 mM Tris pH 8.0, 300 mM NaCl, 20 mM imidazole. gp53 was eluted with 4 column volumes of 50 mM Tris pH 8.0, 300 mM NaCl, 200 mM imidazole. The eluted material was supplemented with 0.5 mM EDTA, 1 mM β-mercaptoethanol and 0.5 mg of the TEV protease, placed in a dialysis bag and left at 20°C overnight for cleavage of the His-tag and dialysis into 20 mM Tris pH 8.0 performed simultaneously. After this step, the protein solution became very viscous. Trypsin was added to this solution to the final amount comprising 1% of the estimated gp53 amount (w/w). The digestion reaction was carried out overnight at 37°C and was stopped on the following morning by addition of PMSF to a final concentration 2 mM. The reaction mixture was concentrated using a Vivaspin 20 ultrafiltration unit with a 5 kDa molecular weight cutoff (Millipore). The concentrated solution was subjected to anion exchange chromatography (GE MonoQ 10/100 GL). The sample was loaded onto the column that was equilibrated with 20 mM Tris pH 8.0 and eluted using a linear 0 to 0.65 M NaCl gradient in 20 mM Tris pH 8.0 buffer. Fractions containing the trypsin digestion product of gp53 (gp53C) were combined and concentrated using a Vivaspin 20 unit. The resulting concentrated solution of gp53C was subjected to size-exclusion chromatography (GE Superdex 200 HiLoad 16/60) equilibrated with 10 mM Tris pH 8.0, 150 mM NaCl. Fractions containing pure gp53C were concentrated using a Vivaspin 20 unit. NaN_3_ was added to a final concentration of 0.02% w/v.

### Crystallization and structure determination

gp53C was crystallized by vapor diffusion at 18°C. Crystals were obtained in approximately 3 days after mixing protein solution at 24.4 mg/ml with crystallization solution that contained 24% PEG 4K, 225 mM Li_2_SO_4_, 100 mM HEPES pH 7.0 in a hanging drop in 1:1 ratio. X-ray data collection was performed using flash-frozen crystals with a cryosolution that contained 50% PEG 4K, 100 mM HEPES pH 7.7, 14% glycerol, 1 M NaBr. Crystals were soaked for ~1 min in this solution to obtain a Br derivative. Diffraction data were collected at the Swiss Light Source (Paul Scherrer Institute, Villigen, Switzerland) at the PX-III (X06DA) beamline using MAR CCD detector. The diffraction data were indexed, integrated, and reduced by the XDS program [13]. Crystals of gp53C belonged to the H32 space group (a = b = 51.20 Å, c = 302.03 Å) with 1 molecule in the asymmetric unit (S1 Table).

The structure of gp53C was solved using the single-wavelength anomalous dispersion (SAD) technique employing anomalous scattering signal from bromide ions. The coordinates of the heavy atom sites (14 per asymmetric unit) and initial phases were obtained using the SHELX pipeline [14,15]. An essentially complete model was then built by Buccaneer [16] and then completed by ARP/wARP [17,18]. It was further refined with PHENIX [11] with manual corrections performed in COOT [12,13]. CCP4 suite of programs was used for various tasks involving the X-ray data and atomic model [22]. The final model contains 20 bromide ions that were placed in peaks higher than 5σ in the Bijvoet difference map. The refinement converged at R-factor of 10% and free R-factor of 13%, and the model was deposited to Protein Data Bank (PDB) under the accession number 4MTM.

### Phage and bacterial stocks

Lytic phage AP22 and its bacterial host-strain *A. baumannii* 1053 were obtained from the State Collection of Pathogenic Microorganisms and Cell Cultures “SCPM-Obolensk” (accession numbers Ph- 42 and B7129, respectively). Phage AP22 was propagated in liquid culture of *A. baumannii* 1053 grown to optical density (OD_600_) of 0.3 (measured at the wavelength of 600 nm) at multiplicity of infection 0.1. Bacterial culture was incubated until complete lysis.

### Lawn spot assay

The enzymatic activity of gp53C was assayed by spotting protein solutions onto the bacterial lawn prepared using the double layer method [23]. 400 μl of *A. baumannii* 1053 bacterial culture grown in LB medium at 37°C to OD_600_ 0.3 was mixed with 4 ml of soft agar (LB broth supplemented with 0.6% agarose). Mixture was plated onto the Nutrient agar (Himedia Laboratories Pvt. Limited, India). 10 μl of the AP22 phage (~109 PFU/ml) and 10 μl of tail fiber gp53C, 10 μl of tail spike gp54 and 10 μl of bovine serum albumin (at concentration 5 μM) were spotted on the soft agar lawn and incubated overnight at 37°C.

### AP22 plaque inhibition assay

In both sets of experiments described below, first, culture of *A. baumannii* 1053 was grown till OD_600_ reached 0.3 in LB medium at 37°C.

In the first set of experiments, 400 μl of *A. baumannii* 1053 culture was mixed with 20 μM of gp53C and incubated for 1 hour at 4°C or for 20 min at 37°C. After the incubation, several dilutions of phage AP22 were added to these mixtures. These solutions were then mixed with 4 ml of soft agar and plated onto the nutrient agar. The plates were incubated overnight at 37°C and assayed for the number of lysis plaques. In a negative control, gp53C was replaced with bovine serum albumin (BSA) at the same concentration.

In the second set of experiments, experimental procedures were identical to those of first set of experiments except *A. baumannii* 1053 cell culture was mixed either with 5 μM gp53C and 5 μM gp54 separately or both proteins together at 5 μM concentrations.

### Bioinformatics

Putative virion proteins of AP22 were identified with the help of protein function and protein structure prediction server HHpred [24]. This tool was used to search against PDB [25], Protein domain families database (Pfam) [26] and Clusters of Orthologous Genes (COGs) [27] databases. Structural homologs of gp53C were found using DALI search (University of Helsinki, Finland) [14] of PDB. Orthologs and of gp53 were identified using HMMER search [29] with the default parameters in UniProtKB database [30]. gp53 sequence was aligned with the sequences of its orthologs using global alignment building strategy in MAFFT 6.0 program [18]. Figure 3B was made with ESPript program [20,21].

### Molecular graphics

UCSF Chimera was used to prepare Figures 2B, 3A and 3D [19]. The electrostatic potential shown in Figure 3A was calculated using the APBS plugin [35] of Pymol [36–38]. Figure 3C was prepared using COOT [20,21].

## Results

### AP22 particle composition

Bioinformatic analysis shows that AP22 is a typical representative of Myoviridae family (Fig 1, Table 1) with virion proteins encoded by a cluster of genes 17 through 54. The arrangement of tail genes resembles that of phage Mu in which neighboring genes encode proteins that interact with each other. Particularly, all the baseplate genes are delineated and arranged by distance of corresponding proteins from the tail axis (Table 1).

**Fig 1.**
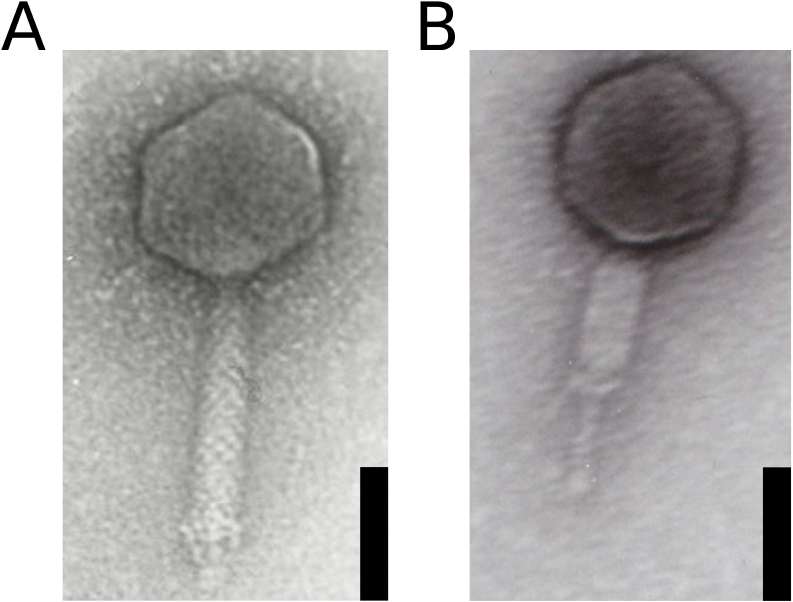
Electron micrographs of CsCl-purified bacteriophage AP22. (Reprinted from [7], with permission from Federation of European Microbiological Societies) (A) Intact phage particle. (B) Phage particle with contracted tail. Staining with 1% uranyl acetate. The scale bar is 50 nm.

**Table 1.**
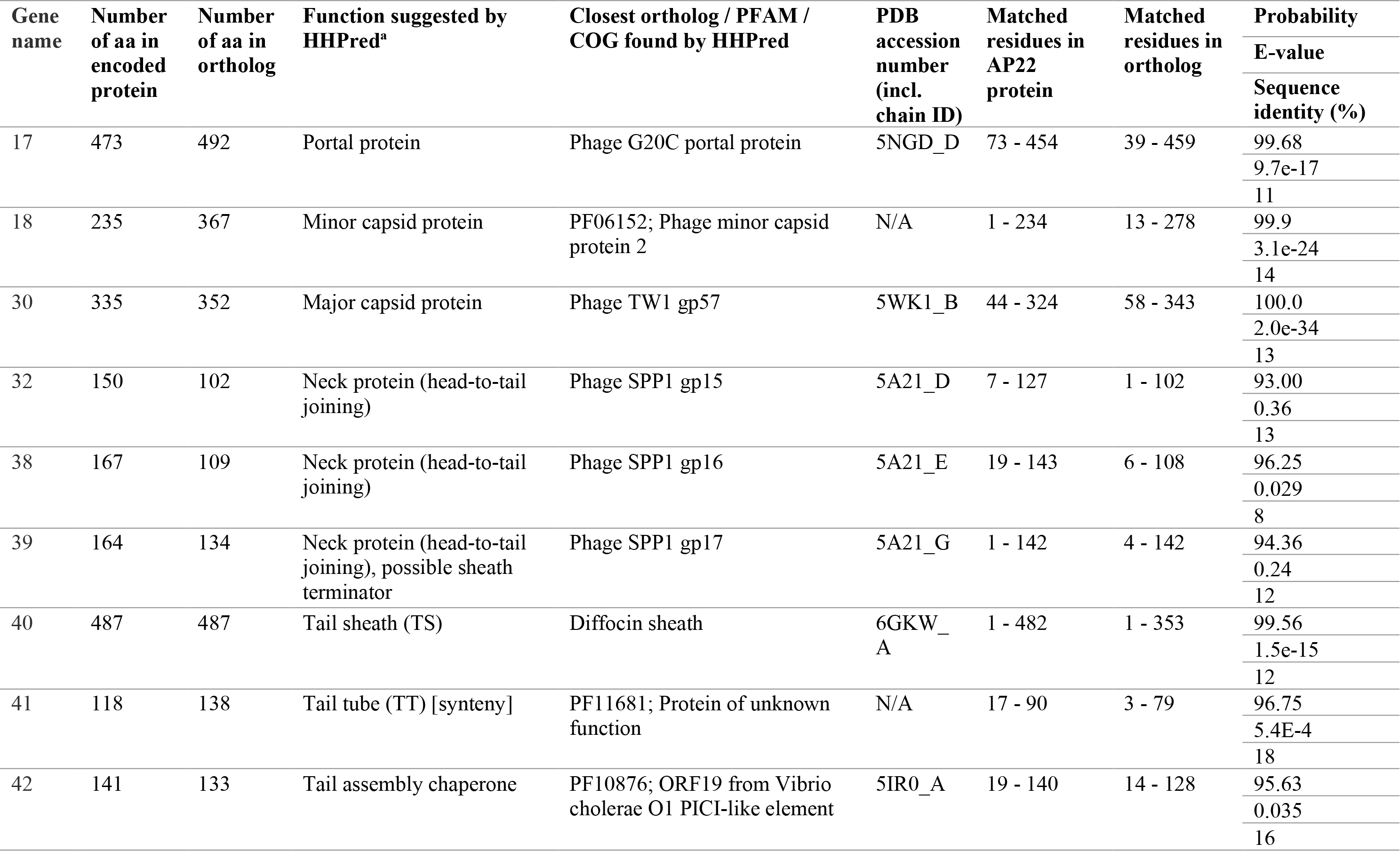
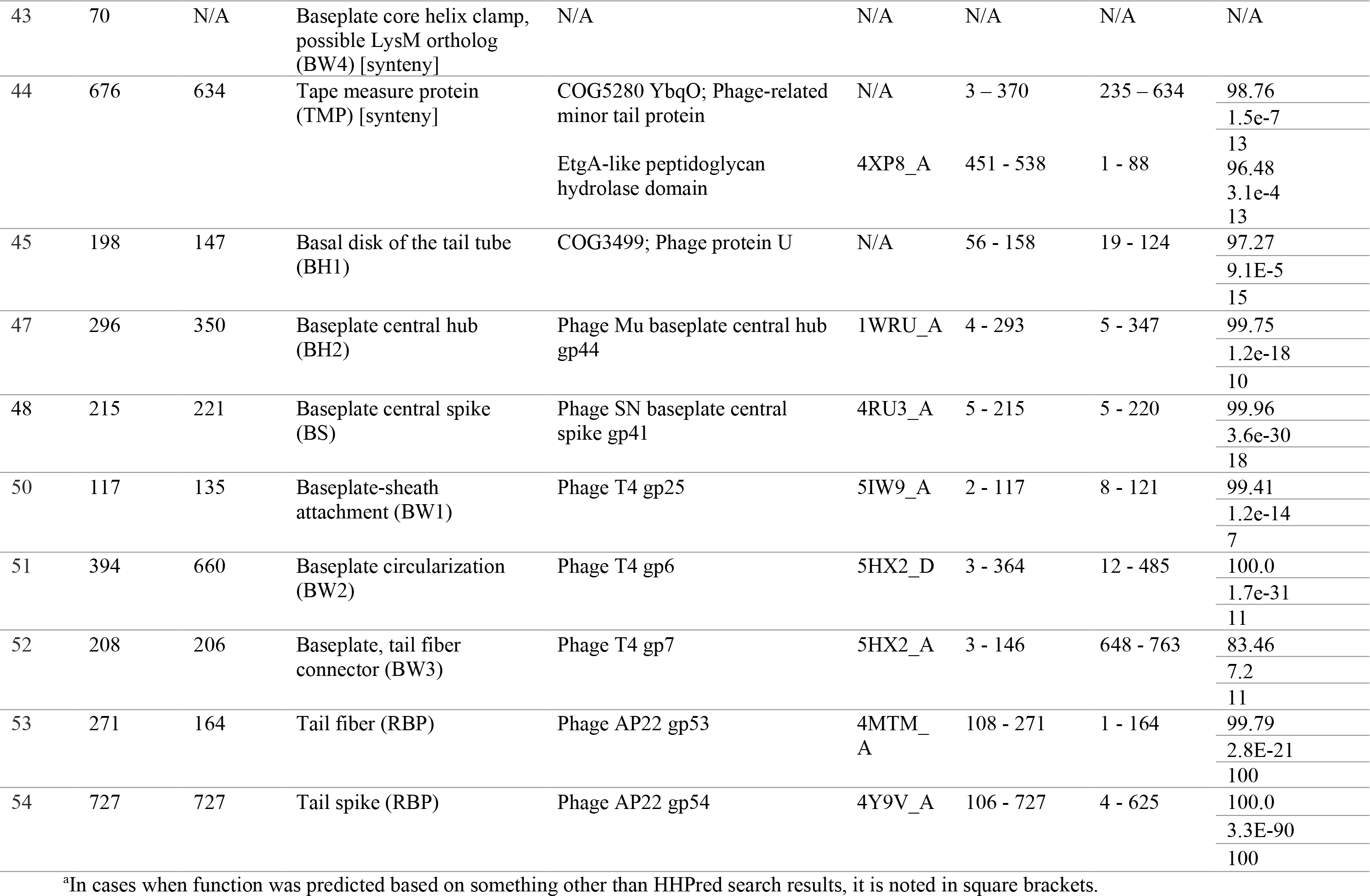
Structural proteins of bacteriophage AP22.

The putative receptor binding proteins are encoded by genes 53 and 54 located at the 3’ end of the virion genome. HHpred analysis shows that a 64-residue fragment of gp53 matches phage T7 gp17 tail fiber with a 98.7% probability and E- and P-values of 4.4 × 10^−9^ and 1.1 × 10^−13^ respectively, and a 414- residue fragment of gp54 matches phage HK620 RBP tail spike with a 99.4% probability and E- and P-values of 2.9 × 10^−10^ and 7.5 × 10^−15^ respectively. SDS PAGE and mass spectrometry show that both gp53 and gp54 are present in the complete phage particle (S1 Fig.).

### Crystal structure of the C-terminal fragment of gp53

Purified full-length gp53 formed a gel-like substance, which lost viscosity upon trypsin treatment as described in Methods. The resulting proteolytic fragment – gp53C (residues Ile135 - Tyr271) – was further purified and crystallized. gp53C is a trimer that shows an unusual mobility on a Superdex 200 10/300 GL gel filtration column most likely due to its elongated shape [39] (Fig 2A). gp53C is a club-shaped molecule in which the “handle” is formed by residues 135 – 172 and the “head” is formed by residues 173 – 271 (Figs 2B, 2C). The handle is a coiled coil domain with an N-terminal lasso-like extension and the C-terminal head is all-beta with lectin-binding fold (Fig 2B).

**Fig 2.**
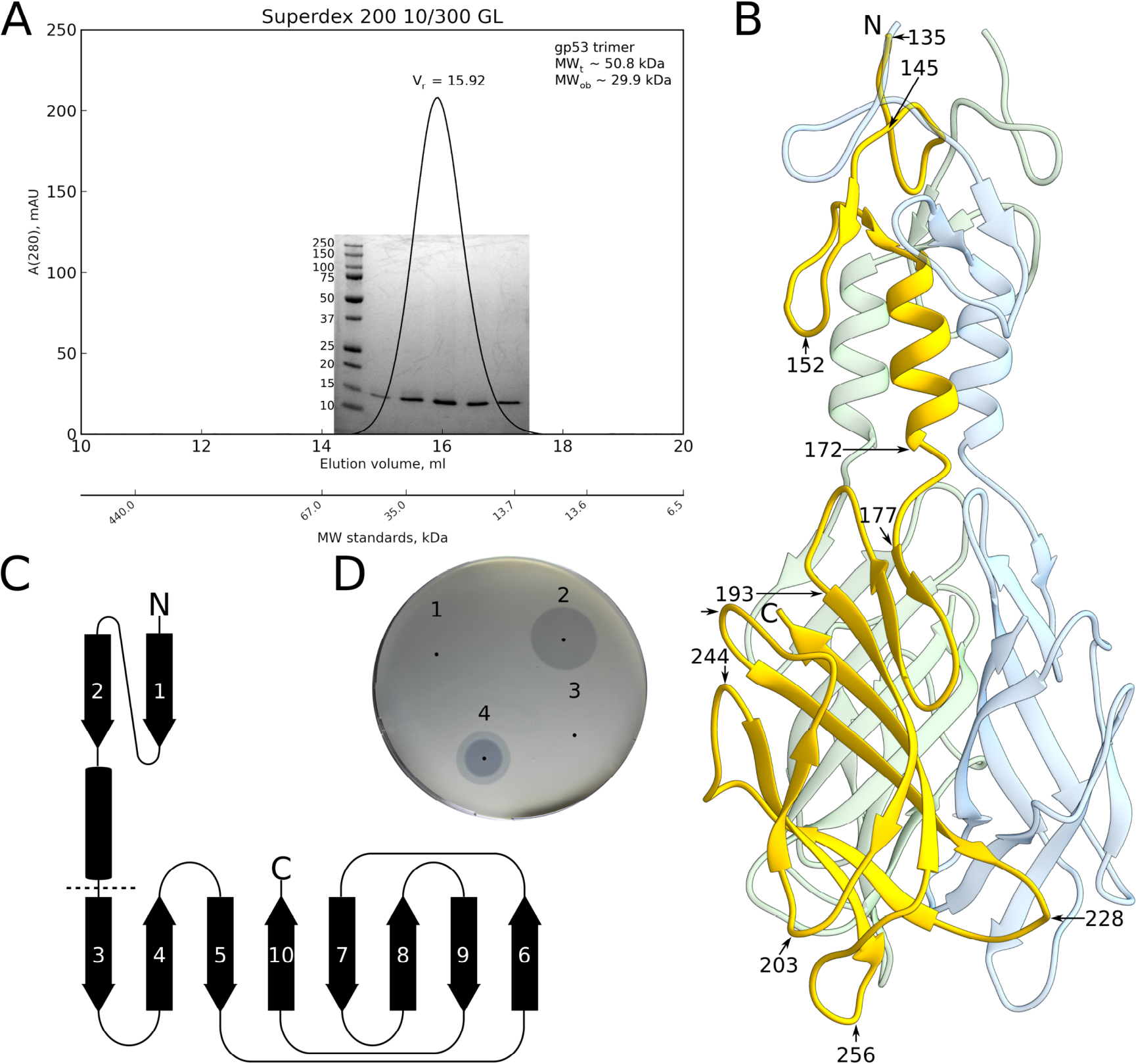
Oligomeric state of gp53C. (A) Size-exclusion chromatogram with overlaid reducing SDS PAGE of the corresponding peak fractions. V_r_ denotes retention volum, MW_t_ denotes theoretical molecular weight, MW_ob_ denotes observed molecular weight. Molecular weights in kDa of protein ladder are shown on the left of SDS PAGE. (B) Crystal structure of gp53C with each chain colored in a unique color. (C) Topology diagram of gp53C monomer. The dash line separates the “handle” and “head” domains. (D) Lawn spot assay of *A. baumannii* 1053 with drops of gp53C (spot 1), gp54 (spot 2), bovine serum albumin (spot 3) and phage AP22 (spot 4). Black dots are added to indicate spot centers.

Several ligands were found to bind to the gp53C surface. Each monomer acquired approximately 20 bromide ions and 1 glycerol molecule from the cryoprotectant solution and 2 ethylene glycol molecules from the mother liquor during crystallization (Figs 3A, 3C and 3D). Halides frequently substitute water molecules in the ordered solvent regions and form ion pairs with positively charged groups such as ε-amino group of lysine, δ-guanidyl of arginine and amide groups of asparagine and glutamine [40,41]. As expected, bromides were found surrounded by positively charged side chains of lysines and asparagines as could be seen in the crystal contact between three neighboring gp53C molecules (Fig 3C).

**Fig 3.**
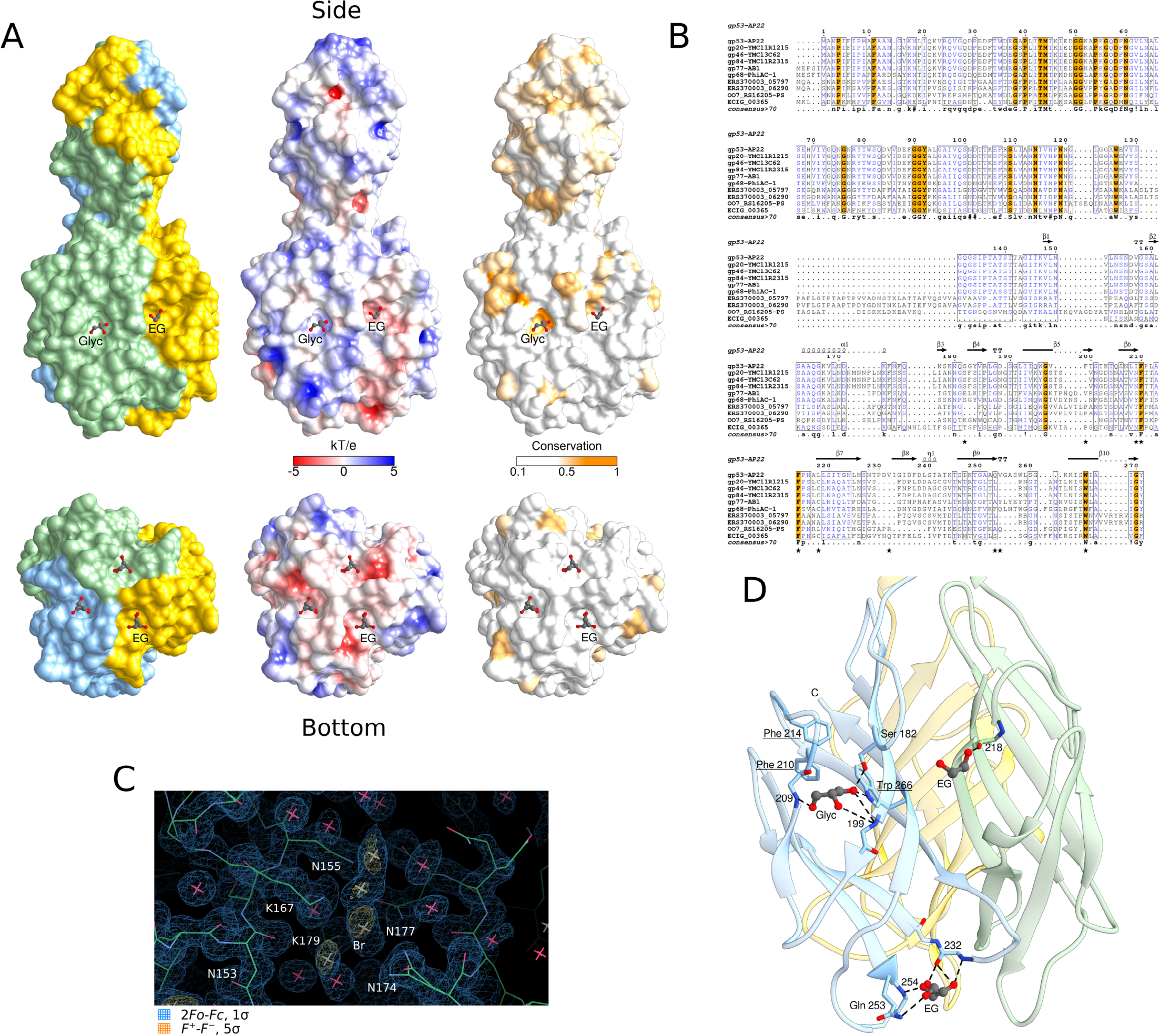
gp53C ligands. (A) Left, Molecular surface of gp53C with each monomer colored in distinct color. Middle, gp53C surface colored by electrostatic potential calculated using Poisson-Boltzmann equation [35]. Right, gp53C surface rendered by conservation based on the alignment of 10 sequences in Fig 3B [47]. Ethylene glycol (EG) and glycerol (Glyc) molecules are shown as ball-and-stick models. (B) MAFFT amino acid sequence alignment of gp53 with its closest homologs: gp20 of *Acinetobacter* phage YMC11/12/R1215, gp46 of *Acinetobacter phage* YMC-13-01-C62, gp84 of *Acinetobacter* phage YMC11/12/R2315, gp77 of *Acinetobacter* phage AB1, gp68 of *Acinetobacter* phage phiAC-1, uncharacterized protein ERS370003_05797 of *Achromobacter* sp., uncharacterized protein ERS370003_06290 of *Achromobacter* sp., hypothetical protein OO7_RS16205-PS of *Providencia sneebia* DSM 19967) and putative prophage protein ECIG_00365 of *Escherichia coli* M605. 100% conserved amino acids are highlighted in orange. Amino acids equivalent in physico-chemical properties are colored blue. Blocks of global similarity are framed. The number in front of each line corresponds to the amino acid position of the corresponding protein sequence. Secondary structure elements – β-sheets, α-helices and turns – are shown as arrows, helices and “TT”, respectively, and are based on the gp53 crystal structure. Stars below the alignment mark the residues involved in coordination of ligands in gp53C structure according to Fig 3D]. The consensus sequence is given for residues that are conserved in 70% or more of all the sequences in alignment. Residues that are conserved in all sequences are shown in capital letters. (C) Anomalous difference Fourier synthesis (F+-F-) contoured at 5σ showing bromide ions located at the site of protein crystal contact. (D) Coordination of one glycerol (Glyc) and two ethylene glycol (EG) molecules in the gp53C structure. Each chain of gp53C is colored in a unique color. Hydrogen bonds are represented by dash lines. Residues that are completely conserved in the set of gp53 orthologs, according to the alignment shown in Fig 3B, are underlined. The glycerol molecule interacts with side chains of S182 and W266 and backbone of I209, F199. F210, F214 and W266 create a hydrophobic part of the glycerol-binding pocket. Ethylene glycol molecule bound on the side of the head domain is coordinated by backbone of L218. Ethylene glycol molecule bound at the tip of gp53C interacts with side chain of Q253 and backbone of V232, V254.

Glycerol and ethylene glycol molecules bound to gp53C are coordinated by hydrogen bonds (Fig 3D). One of the ethylene glycol molecules is in a small cavity at the tip of the head domain whereas the other is in a cavity near the widest part of the head domain. The glycerol molecule occupies a larger cavity roughly in the middle of the head domain. Because a polyethylene glycol chain mimics an oligosaccharide backbone in an extended conformation [42], it is possible that these pockets bind sugar receptors extending from the host cell surface.

### Structural homologs of gp53C

The overall shape of gp53C resembles that of the receptor-binding part of the phage T4 short tail fiber gp12 and, to a lesser degree, the T7 tail fiber gp17. The latter was identified as a possible structural ortholog of gp53 in an HHpred search (Table 1). The matching region of the two proteins is a fragment of the C-terminal globular domain, which presumably is involved in LPS binding in T7 gp17. The overall folds of these domains, however, are different.

The gp53C C-terminal head domain has a lectin fold (an 8-stranded antiparallel beta-barrel) that differs from folds of either T7 gp17 or T4 gp12. However, this fold is fairly ubiquitous among proteins of non-immune origin that bind carbohydrates [43]. A search for structural homologs using the DALI server against the non-redundant subset of PDB yields 141 hits. The top hits that show both low RMSD and high Z-score are agglutinin of snail *Helix pomatia* (PDB ID 2CGZ, RMSD 2.6 Å, Z-score 5.9), RBP of the *Lactococcus lactis* phage bIL170 (PDB ID 2FSD, RMSD 2.9 Å, Z-score 5.7) and endorhamnosidase tail spike of *Shigella flexneri* phage Sf6 (PDB ID 2VBK, RMSD 3.4 Å, Z-score 5.0). Many of these proteins have been implicated [44,45] in binding oligosaccharides or lipopolysaccharides.

Interestingly, there is a similarity in an important element of T4 gp12 and AP22 gp53 proteins. Namely, the very N-terminal part of the gp53C crystal structure is a lasso-like motif that excellently match to the repeating unit of T4 gp12 (Fig 4). The two have low (18%) sequence identity and yet can be superimposed with an RMSD (root-mean-square deviation) of 4.2 Å between 32 aligned C-alpha atoms. Why T4 gp12 and AP22 gp53 proteins have similar structural element despite significant difference in their overall folds, is not clear.

**Fig 4.**
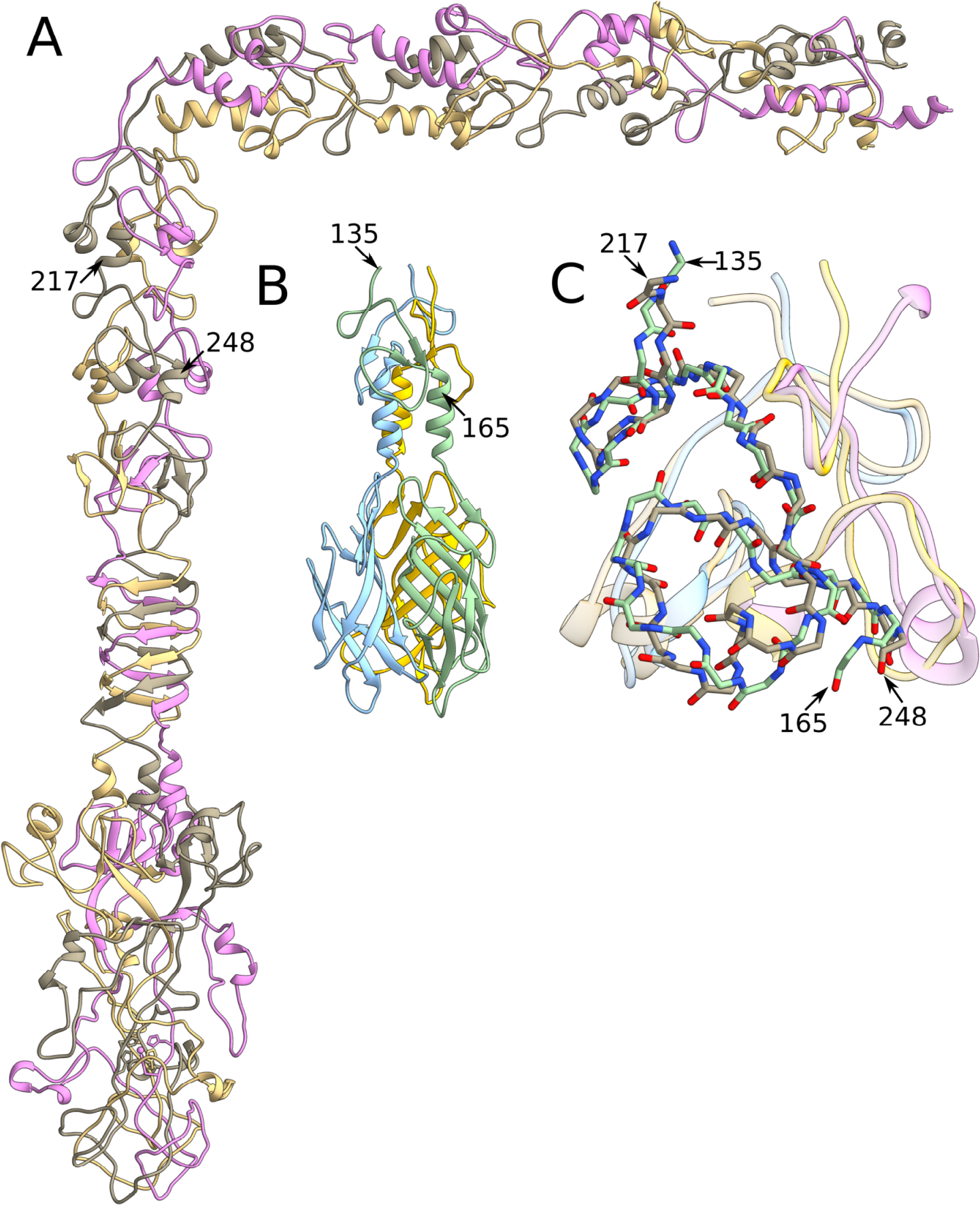
Conservation of the lasso motif of AP22 gp53 to T4 gp12. (A) T4 gp12 structure with each chain colored in a unique color. (B) AP22 gp53 with each chain colored in a unique color. (C) Superposition of T4 gp12 onto AP22 gp53. Two chains in each trimer are shown as a semitransparent ribbon diagram and one monomer is displayed as an all-atom backbone. Structurally similar fragments in the two proteins are indicated with numbered residues.

### gp53-like proteins

Currently, GenBank contains 9 homologs of gp53 (Fig 3B). Their sequences align with gp53 sequence reasonably well and have lengths similar to that of gp53. Their N-terminal parts are conserved, whereas C-terminal regions show greater sequence variability, typical of phages’ RBPs [10,46] (Figs 3A, 3B). We mapped sequence conservation of all ten gp53-like proteins onto the molecular surface of gp53C (Fig 3A). Interestingly, the glycerol binding pocket is likely to be present in all homologs since residues forming this pocket are conserved (Figs 3A, 3B). The ethylene glycol binding pocket on the side of the head domain is semi-conserved (Figs 3A, 3B). Another ethylene glycol binding pocket, at the tip of the head domain, may not be present in all homologs because amino acid residues making up this pocket in gp53 are not conserved (Figs 3A, 3B).

This analysis of amino acid sequences suggests that the binding pocket occupied by the glycerol molecule in gp53C could be a universal feature of all gp53 orthologs. The glycerol molecule interacts with a side chain of S182 and W266, and the latter is one of the few residues universally conserved in all orthologs as well as F210 and F214 that together with W266 create a hydrophobic part of the glycerol-binding pocket (Figs 3B, 3D). Backbones of I209 and F199 amino acid residues are also involved in coordination of glycerol molecule (Fig 3D).

Ethylene glycol binding pocket on the side of the head domain could also be present in all gp53 orthologs, but its shape and properties are likely to differ since only one residue (L219) out of four forming this pocket (Q181, V198, L219 and L239) is somewhat conserved (Figs 3B, 3D). At the same time, L218, that actively participates in coordination of this ethylene glycol molecule in gp53, is not conserved in these gp53 homologs (Figs 3B, 3D). Interestingly, in 5 homologs out of 10 there is cysteine in this position (Fig 3B).

Finally, the ethylene glycol binding pocket at the tip of the structure might be present in some homologs but absent in others. Residues that form this pocket (Q253, V232, V254) in gp53C are not conserved (Figs 3B, 3D).

### Exploring gp53 function

In the lawn spot assay, phage AP22 forms clear lysis zones surrounded by opaque halos that expand with time suggesting that an AP22-encoded enzyme diffuses away from the lysis zone and degrades the thick external polysaccharide of *A. baumannii* cells. We surmised that this phenomenon could be due to one of the proteins used by the phage for attachment to the host cell. We performed the spot assay using gp53C, the other putative RBP gp54, and bovine serum albumin (BSA) as a control using 130 multi-drug resistant *A. baumannii* isolates from hospitalized patients of Chelyabinsk, Nizhny Novgorod, Moscow and St. Petersburg in 2005-2013. The spotting of gp54 resulted in the appearance of halos similar to halos appearing on bacterial lawns of *A. baumannii* strains that previously were shown to be susceptible to AP22 infection. In contrast, the spotting of gp53C in a concentration up to four times higher than that of gp54 did not result in lysis or formation of halos (Fig 2D).

We decided to investigate if either gp53 or gp54 can block AP22 infection by binding to or modifying the surface of an *A. baumannii* cell using plaque inhibition assay. To do so, *A. baumannii* bacteria were first incubated with either gp53C or gp54, then AP22 phage was added and the whole mixture was plated. After an overnight incubation, phage titer was measured. Incubation with gp53C did not affect infection of *A. baumannii* cells with AP22 as the phage titer was very similar to that resulted from infection of *A. baumannii* cells with AP22 without prior incubation of bacteria with anything (Fig 5). This experiment was repeated with varying amounts of added protein and incubation conditions (incubation temperature, length of incubation). However, no significant differences in phage titers were detected. In contrast, co-incubation of *A. baumannii* bacteria with gp54 had a profound effect – after that *A. baumannii* cells became non-susceptible to AP22 infection (Fig 5). These results suggest that gp54 is likely to either bind tightly the *A. baumannii* external polysaccharide capsule that serves as a host cell receptor for AP22 and thus prevent AP22 binding or degrade this capsule and thus prevent AP22 binding.

**Fig 5.**
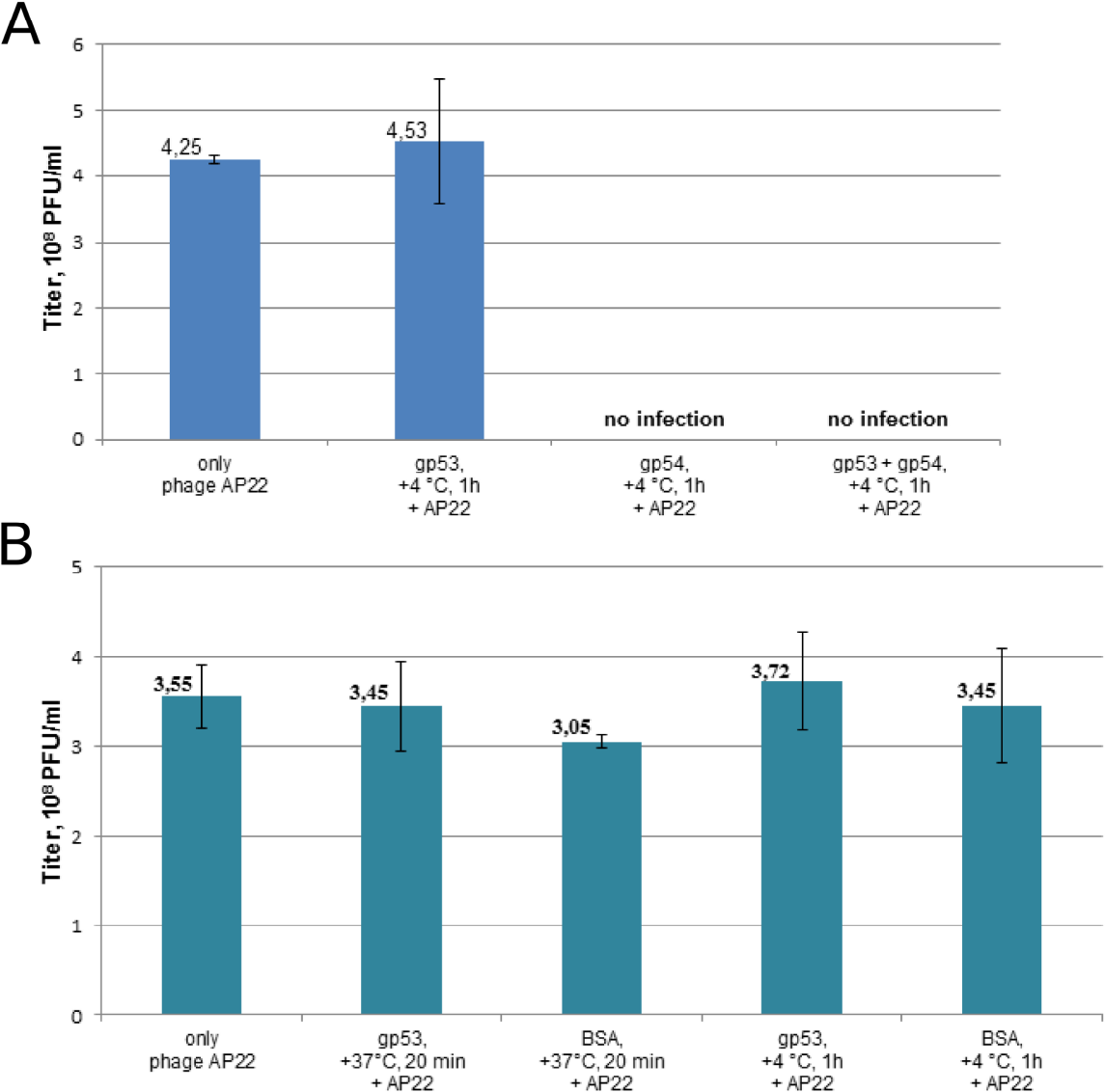
AP22 plaque inhibition assay. (A) Titer of phage AP22 after infection of *A. baumannii* 1053 cells that were incubated with either gp53C or gp54 or gp53C-gp54 mixture. *A. baumannii* cells not incubated with anything served as control. Error bars are standard deviations for two different phage dilutions tested. (B) Titer of phage AP22 after infection of *A. baumannii* 1053 cells incubated with gp53C at 37°C for 20 minutes or at 4°C for 1 hour. *A. baumannii* 1053 cells not incubated with anything and cells incubated with BSA served as controls. Error bars are standard deviations for two different phage dilutions tested.

## Discussion

We determined the crystal structure of the C-terminal fragment of phage AP22 gene product 53 that comprises approximately 50% of the total sequence length. We found that the protein is a structural component of the phage particle and displays several features that are common to phage RBPs starting from the overall sequence conservation (conserved N-terminus and divergent C-terminus) and overall club-like shape of the molecule and continuing with a C-terminal lectin domain, shared motifs with T4 short tail fiber gp12, conserved pockets on the lectin domain that bind ethylene glycol and glycerol. However, we were not able to identify the exact function of gp53. It does not display enzymatic activity and does not appear to modify the surface of *A. baumannii* in a way that was detectable in our functional assays. Furthermore, it does not appear to inhibit phage infection. It is possible that gp53 binds to a receptor on the cell surface that is hidden by the capsular polysaccharide so that receptor’s uncovering requires a spatially and temporarily defined action of gp54 or some other AP22 tail protein.

## Acknowledgments

We sincerely thank the staff of the protein crystallography beam lines at the Swiss Light Source and especially Vincent Olieric, Meitian Wang, Takashi Tomizaki, Martin Fuchs, Florian Dworkowski and Clemens Schulze-Briese for their help with X-ray data collection and support.

## Supporting Information

**S1 Table. gp53C data collection and refinement statistics.**

**S1 Fig. Structural components of AP22 virion.** (A) SDS PAGE of complete AP22 particle. Molecular weights in kDa of protein ladder are shown on the right of SDS PAGE. (B) Mass spectrometry analysis of the trypsin digestion products of the protein extracted from the SDS PAGE labeled as gp53. Twelve large peptides uniquely identify gp53.

