## Supplementary material for "Crystal Structure of the putative tail fiber protein gp53 from the *Acinetobacter baumannii* bacteriophage AP22": S1 Table, S1 Fig

**S1 Table.** **gp53C data collection and refinement statistics.**

| Data collection | Br f″ peak |
| --- | --- |
| X-ray source | SLS, PX-III |
| Wavelength (Å) | 0.9200 |
| Space group | H32 |
| Unit cell (Å) | 51.20, 51.20, 302.03 |
| Resolution range (Å) | 19.72 - 1.37 |
| Unique reflections | 31897 (2865)* |
| Multiplicity | 10.6 |
| Completeness (%) | 96.49 (88.43) |
| <I>/σ(I) | 24.16 (3.69) |
| Wilson B-factor (Å^2^) | 8.87 |
| SigAno (\|F^+^-F^-^\|/σ) | 1.262 |
| Refinement |  |
| R (%) | 9.93 (14.90) |
| R_free_ (%) | 13.17 (20.15) |
| Number of atoms: |  |
| macromolecule/ligand/water | 1049/43/149 |
| RMSD of bond lengths (Å) | 0.01 |
| RMSD of bond angles (°) | 1.32 |
| Ramachandran favored residues (%) | 97 |
| Ramachandran outliers (%) | 0 |
| B-factors (Å^2^): |  |
| average/macromolecule/solvent | 12.80/10.30/25.70 |
| *The statistics in the parenthesis is for the highest resolution range bin of 1.42-1.37 Å. | |

**S1 Fig.** **Structural components of AP22 virion.** (A) SDS PAGE of complete AP22 particle. Molecular weights in kDa of protein ladder are shown on the right of SDS PAGE. (B) Mass spectrometry analysis of the trypsin digestion products of the protein extracted from the SDS PAGE labeled as gp53. Twelve large peptides uniquely identify gp53.

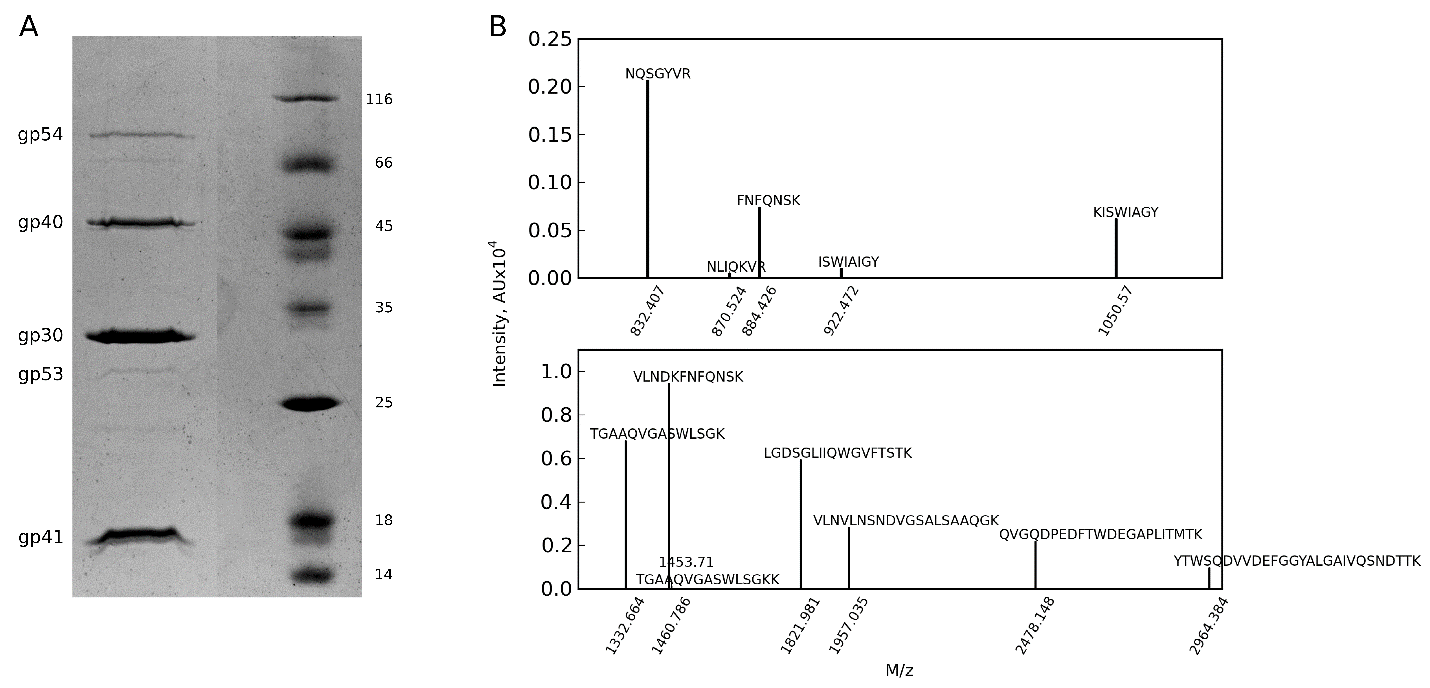
